# Small hand-designed convolutional neural networks outperform transfer learning in automated cell shape detection in confluent tissues

**DOI:** 10.1101/2022.10.17.512515

**Authors:** L. Combe, M. Durande, H. Delanoë-Ayari, O. Cochet-Escartin

**Affiliations:** Institut Lumière Matière, UMR5306, Université Lyon 1-CNRS, Université de Lyon, 69622 Villeurbanne, France; Laboratoire Matière et Systèmes Complexes, UMR7057, Université Paris Cité-CNRS, F75205 Paris Cedex 13, France

## Abstract

Mechanical cues such as stresses and strains are now recognized as essential regulators in many biological processes such as cell division, gene expression or morphogenesis. Studying the interplay between these mechanical cues and biological responses requires experimental tools to measure these cues. In the context of large scale tissues, this can be achieved by segmenting individual cells to extract their shapes and deformations which in turn inform on their mechanical environment. Historically, this has been done by segmentation methods which are well known to be time consuming and error prone. In this context however, one doesn’t necessarily require a cell-level description and a coarse grained approach can be more efficient while using tools different than segmentation.

The advent of machine learning and deep neural networks has revolutionized the field of image analysis in recent years, including in biomedical research. With the democratization of these techniques, more and more researchers are trying to apply them to their own biological systems. In this paper, we tackle a problem of cell shape measurement thanks to a large annotated dataset. We develop simple CNNs which we thoroughly optimize in terms of architecture and complexity to question construction rules usually applied. We find that increasing the complexity of the networks rapidly no longer yields improvements in performance and that the number of kernels in each convolutional layer is the most important parameter to achieve good results. In addition, we compare our step-by-step approach with transfer learning and find that our simple, optimized CNNs give better predictions, are faster in training and analysis and don’t require more technical knowledge to be implemented. Overall, we offer a rational roadmap to develop optimized models and argue that we should limit the complexity of such models. We conclude by illustrating this strategy on a similar problem and dataset.

## Introduction

It is becoming abundantly clear that the mechanical environment encountered by cells, tissues and organisms plays an important regulatory role intertwined with chemical or metabolic regulations. At the cell level, mechanical cues can influence processes as important as cell division (1,2), individual and collective migrations (3–5), gene expression (6,7) or even cell fate (8,9). These discoveries are at the root of the rapid expansion of the mechanobiology field. In this context, it is crucial to develop tools to quantitatively characterize the mechanical environment encountered by cells *in vivo* or in *in vitro* experiments. Experimental tools are being developed to measure relevant mechanical properties *in situ* such as deformable force sensors (10,11), FRET-based mechanical reporters (12,13) or other optical techniques to measure cell membrane tension (14).

In the case of living tissues, it has long been recognized that mechanical information can be inferred from cell shapes, deformations and how they change over time (15–18). Notably, observation of cell shapes can even be predictive as to the mechanical behavior of the tissue as a whole (19). The most natural way to estimate cell shapes is to automatically detect their contours and therefore segment them from one another giving access to shapes and deformations on a single cell basis. These segmentation techniques can be manual, automated by usual image analysis tools or automated by machine learning based techniques. Still, these techniques are notoriously error-prone, often requiring manual corrections. More problematic, small errors in segmentation will tend to fuse neighboring cells together thus having a large impact on the desired measurements. In many cases, these techniques can still be applied with a relative efficiency to yield cell level data.

However, in the context of large scale tissues, cell-level information might not be required and a coarse grained representation can be sufficient. Relevant mechanical data can be obtained by estimating the local inertia tensor, the equivalent of measuring an average cell size, locally. Such approaches can be achieved even in the absence of cell segmentation, for instance by using 2d Fourier transforms (20).

In the field of biomedical image analysis, many recent advances have been achieved by the application of machine learning and, in particular, of convolutional neural networks (CNNs). Historically developed to treat everyday images and perform tasks such as object detection or classification, the use of these algorithms has now penetrated deep into biological research. Examples of application of CNNs to microscopy images are rapidly being added in contexts as different as crowded cell environments (21), single cell classification (22,23), neuronal imaging (24) and of course cell segmentation (25–27). These models have thus already demonstrated their strengths in aiding biological images analysis, in large part thanks to the increasing availability of GPUs and computing power in general. Still, CNNs are complex mathematical models and they can easily suffer from pitfalls and biases (28) if not trained properly or if the dataset used is not properly designed.

As a result, in many instances, researchers will not try to develop their own neural networks and optimize them for a specific problem. Instead, they use an approach called transfer learning. The underlying idea is to make use of already designed and already trained neural networks and re-train them to tackle a new problem. Powerful models are capable, thanks to their training, complexity and underlying dataset, of creating very detailed representations of an image in order to extract information. The idea of transfer learning is to freeze the first layers of such pre-trained models, arguing that the learned representations will also be relevant for the problem at hand. The last layers of the model, the ones which actually make a prediction from an image, are then either re-trained or replaced as the output of the problem is often very different from the one they were originally trained on. This technique has proven quite powerful in a variety of cases (29–31).

However, deep and transfer learning are complex tools and some of the drawbacks of their intensive use have started to appear. These range from the use of deep learning where it is clearly neither adapted nor required (32,33) through its weaknesses to adversarial attacks (34), the reproduction of biases and to examples where seemingly efficient models fail to generalize to new, albeit very similar situations due to domain shift (35,36). One should thus be careful in applying deep learning techniques, explore their own data and optimize models to fit specific needs. This might seem like a complicated task that can be overcome by the use of transfer learning. However, it is unclear in which situations transfer learning is indeed a good alternative and if it is actually simpler than developing one’s own CNNs from scratch. Based on a real-life example in biophysics, we will explore this question.

In this paper, we made use of a large scale, high quality annotated data set coming from (20) and experimentally obtained in (37). It contains large, raw images of confluent fly dorsal thorax tissue images. Thanks to a partially manual segmentation of these images, it also contains the exact shape of all cells within the tissue, allowing for the definition of a ground truth. We started by properly defining our coarse grained cell shape detection problem. Then, we created and explored a dataset designed to address this problem. To solve it, we first developed our own CNNs from scratch and started from the simplest possible ones. We then increased complexity step by step in order to optimize our model without using overcomplicated ones. In this step, we also extensively explored the underlying hyperparameters to test the architecture rules of thumbs which are usually applied in building CNNs but which are not grounded in hard data. We also tested how the increasing complexity of the network increased its performances. We ended up with a fully optimized, sparse, 3-layer model which achieved great performance although it diverged from usually applied architecture rules. Finally, we compared our homemade, step-by-step approach with a more direct transfer learning strategy and showed how the former outperforms the latter in terms of performance, required expertise and computing time. We also test the influence of the size of the data set on both approaches and find that our results hold true even for small training sets.

Overall, we 1-offer a very robust algorithm for cell shape detection in confluent tissues at a coarse grained level, thus offering a biologically relevant image analysis tool and 2-through extensive exploration of both data and models, we question common practices encountered in similar situations and offer a rational, optimized way of designing efficient CNNs to solve relatively simple image analysis problems without the need to use transfer learning. We finally illustrate these conclusions on a different data set and show how simple CNNs can achieve a high level of performance in less than a day in cell measurement image analysis.

## Results

### Definition of the problem and data exploration

The aim of this work was to create a model capable of producing a coarse-grained map of inertia tensors over a large biological tissue from images only. We started from images of a single fly dorsal thorax tissue (Fig 1A). We cut this image in multiple smaller images giving the resolution of our coarse grained level. Inputs to the models were black and white images 128*128 pixels in size involving a few tens of cells (Fig 1B). Targets (ground truth) were related to averaged inertia matrices giving an average cell shape. Our ground truth was calculated thanks to semi-automated segmentation (Fig 1B). This shape is fully characterized by 3 parameters: length, width and orientation (Fig 1B) of an average cell in the corresponding region. We therefore faced an image analysis, multiple regression problem. Thanks to previous work, we started from a large dataset of 17146 images separated into a training set (15989 images) and a test set (1157 images).

**Fig1.**
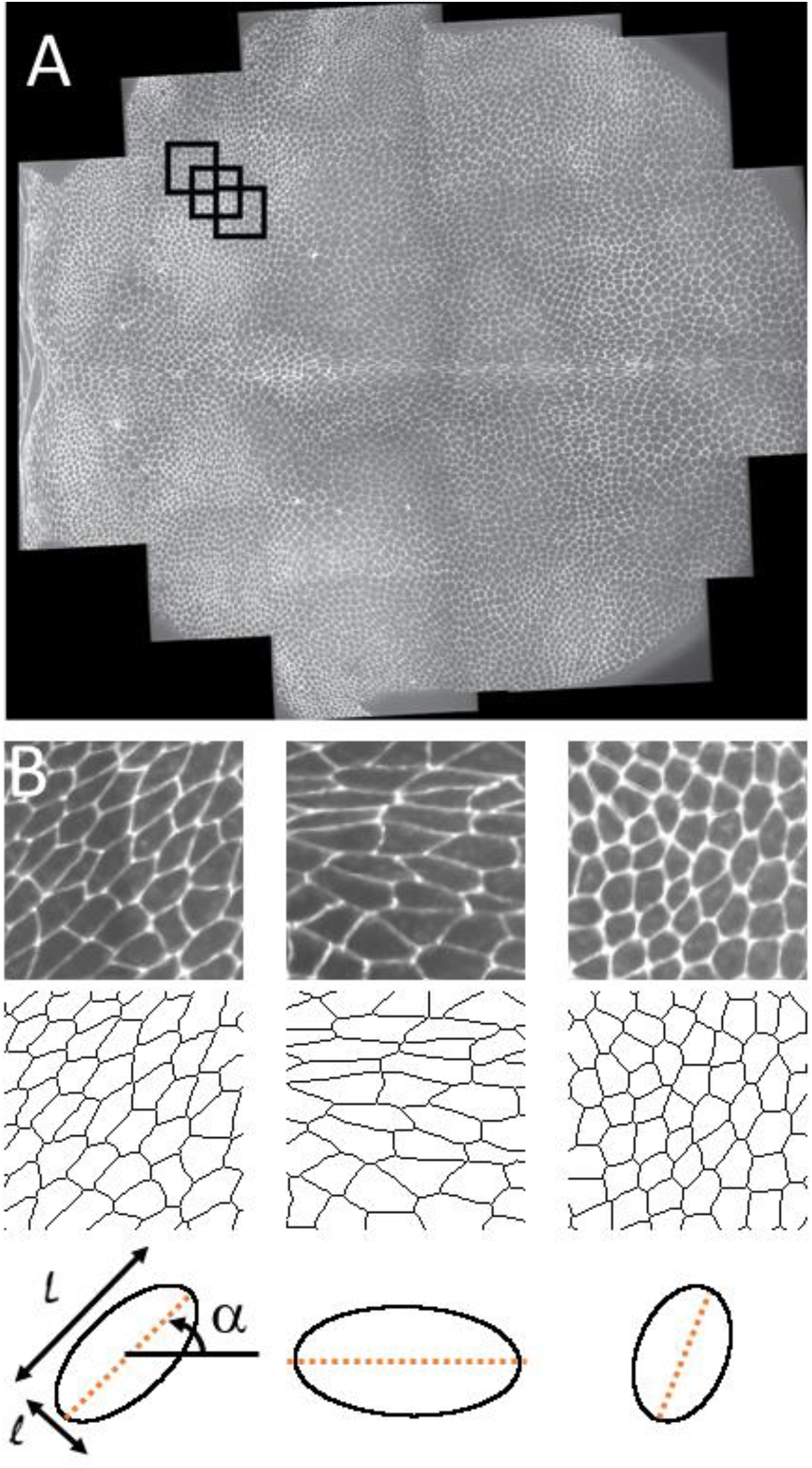
Illustration of the problem. A: full raw image of the Drosophila dorsal thorax. Small black circles represent the size of the images used in our models. B: three examples of such 128*128 images along with their segmented counterparts and representations of the average cell shapes demonstrating the three key targets: long axis L, short axis l and orientation α (angle between the horizontal and the orange line).

Before defining any model or training strategy, we started by exploring our annotated data both in terms of the inputs and the targets. The inputs were simple enough but we still looked at the distribution of grey levels over all images in the dataset to see if we detected any outliers or problematic values (Fig 2A). We found that the distribution seemed coherent but that grey values were spread between 0 and 255. As common practice, we decided to rescale all images by dividing them by 255, ensuring that all inputs were bound between 0 and 1 while avoiding artefacts potentially stemming from rescaling each image from its maximum.

**Fig 2.**
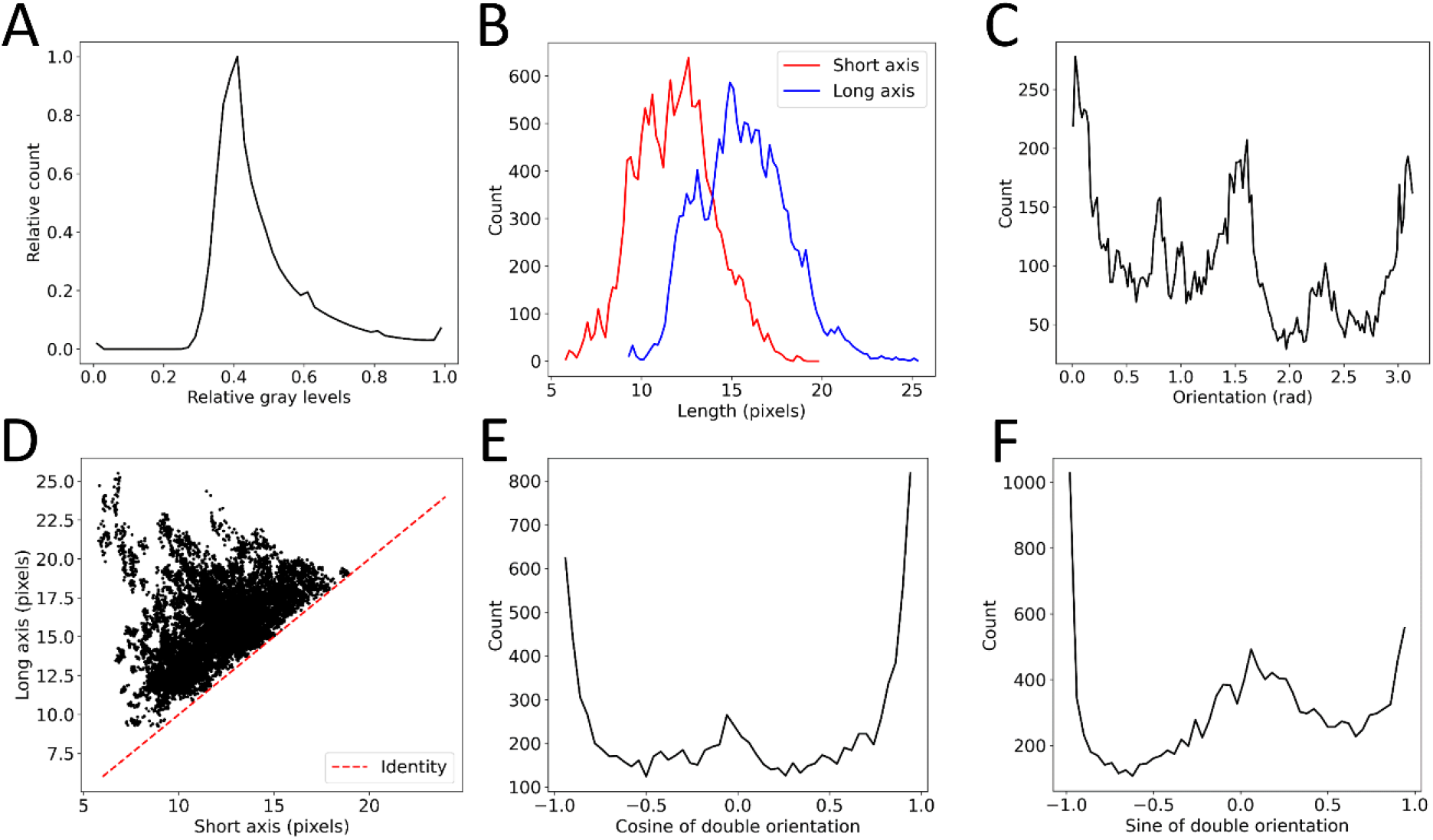
Exploration of the data set. A: relative gray level distribution over the entire dataset. B: Distributions of long axis, in blue, and short axis, in red, in the training set. C: distribution of the raw orientation in the training set. D: Relationship between short and long axes in the training set, the red dashed line showing y=x to demonstrate the correlation between the two variables. E: Distribution of the cosine of double the orientation. F: Distribution of the sine of double the orientation.

Regarding the targets, we also looked at their original distributions to detect and remove outliers, if any. Thankfully, the data set was remarkably clean and no clear outlier emerged (Fig 2B).

Interestingly, the angular distribution was not homogenous and values of 0, π/2 and π seemed overrepresented (Fig 2C). In addition, we did find some structure in the data. Obviously, the long and short axes of the averaged cells were somewhat correlated since they had to obey short axis < long axis (Fig 2D). Apart from that, they didn’t seem to be correlated enough for us to remove one and infer it simply from the other. We thus kept them both as two independent targets while keeping in mind that the data is partially structured.

Another conclusion was that orientation was a more problematic quantity. It is, by definition, bound between 0 and π, it is periodic, *i*.*e*. values of 0 and π actually represent the same measurement and it has a non-normal distribution. As a result, usual loss functions such as mean squared or absolute errors seemed ill-suited for this particular target as they would heavily penalize predictions close to π for a ground truth close to 0, and vice versa. This raised the question of how to encode the orientation data and which loss function to use.

We tested two different possibilities: keep the orientation encoded as angles between 0 and π but use a periodic mean squared error (MSE) as a loss function or encoding them as two targets: the cosine and sine of the double of the angle (Fig 2E-F) using a regular MSE as a loss function. Doing so, we ensured that the target was first resampled on [0, 2 π [and then that the (cos, sin) encoding was no longer periodic. Unlike the original encoding, large differences in cosine and sine couldn’t come from very similar data.

Empirically, the latter seemed to work better (Fig SI1) and we kept this approach as it yielded better performance, was more natural and user friendly (no need to manually define loss functions). It might seem counterproductive as we created an additional target, seemingly increasing the complexity of the problem, and adding correlations between these targets. Also, we had to define an inverse transformation which is non-unique. However, we found from a simple convolutional network example that these CNNs can easily learn basic geometric laws as the correlation between predicted cosines and sines closely followed the rule cos^2^+sin^2^=1 (Fig SI3). This result also showed that a 2-argument arctangent was a good way to perform the inverse transformation and obtain the orientation from the couple of values (cos,sin).

From then on, we focused on this new definition of the problem: from binary images of 128*128 pixels, we wanted to develop models predicting the average inertia matrix encoded as length, width, cosine and sine (of twice the orientation) using a usual MSE as a loss function.

Another question we investigated was whether a single model is capable of fitting both lengths and angles or if it was more pertinent to separate lengths and orientations in our approach. We compared performances of very simple 2d CNNs in each situation. By fitting all targets at the same time, we noticed a very small loss in performance (Fig SI2) but this approach was much more practical. The fact that a single model was capable of fitting data of different nature was not a given and indicated that our approach which aims at predicting complex quantities with simple CNNs was relevant.

We now had a well-defined regression from images problem with four targets, a dataset that corresponded to this problem and had a both a large number of images and high quality ground truth. It was therefore perfectly suited to the use of supervised learning and CNNs in particular. Following our strategy, we started by developing the simplest possible CNNs and increased complexity step by step until achieving acceptable performance on our problem. Our aim was thus to compare different models with one another and select the best performing one. As is usual practice, we used a 5-fold cross validation strategy. Each of our models was thus trained 5 times in such a way that each image in the training set was used four times as actual training and once as validation. The performance of a given model was given as the mean absolute error (MAE) on the validation set, averaged over the 5 folds.

In terms of architecture, our CNNs were based on the repetition of the same module consisting of a convolutional layer with Relu activation, a 2 by 2 maximum pooling and a dropout layer set at 0.2 (Fig3). After N repetitions of this module, we simply flattened the output of the last convolutional layer and added a 4 neuron dense layer with linear activation as the final regressor outputing long axis, short axis, cosine and sine of the orientation.

**Fig3.**
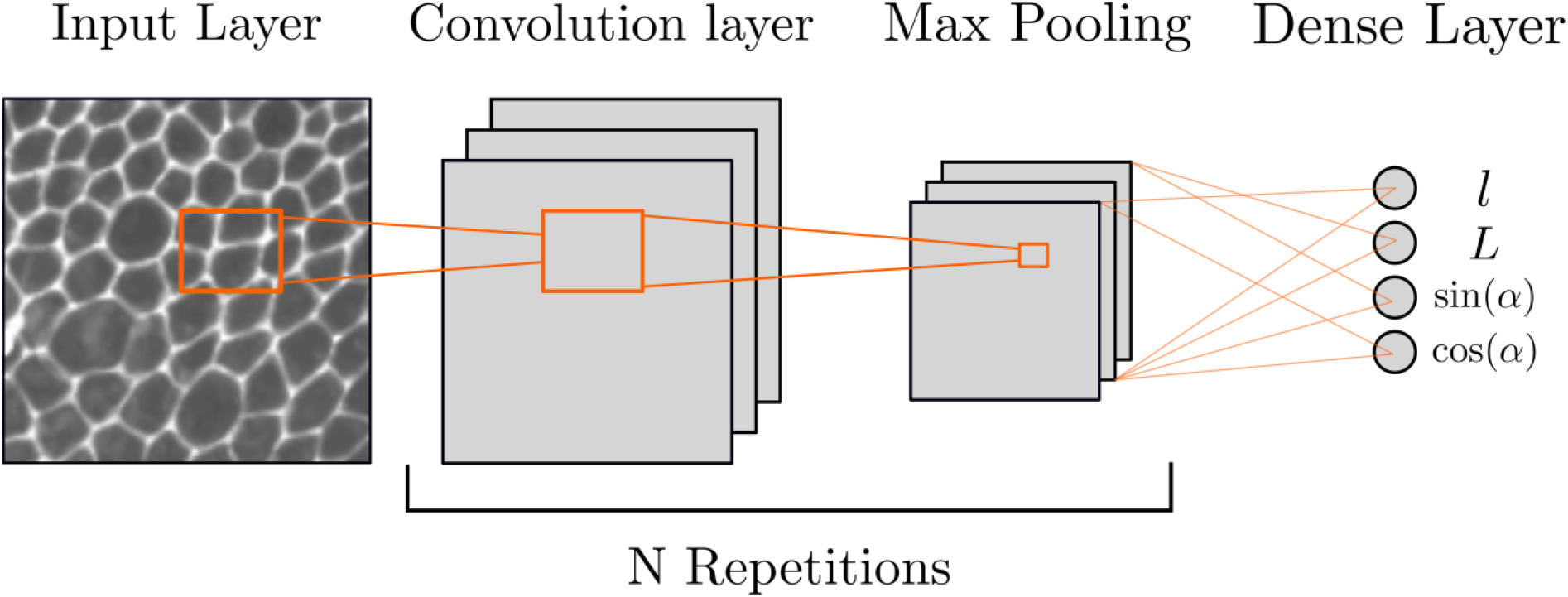
Schematics of the networks’ architecture. Models are based on the repetition of convolutional and max pooling layers. The dropout layers are not represented to help visualization. The four neurons of the final dense layer each correspond to one of the model’s targets, L being the long axis, l the short axis and α the orientation.

In this work, we extensively studied the hyperparameters of the convolutional layer: the number and size of convolutional filters, or kernels, to be learned by the model. These hyperparameters can have a large impact on the final performance of the model but optimal values are hard to predict. Some rules of thumbs exist and usually are as follows. The number of kernels is a power of 2 (4,8,16 etc..) and is multiplied by 2 for each new convolutional layer. The size of the kernels, in pixels, is kept constant. Thanks to the maximum pooling, this means that kernels in each new convolutional layer actually treat a larger area of the original image, effectively zooming out.

### Optimized 1 convolutional layer model

As described, we started by the simplest possible case, corresponding to N=1 repetition. We independently varied the numbers and sizes of the kernels in this convolutional layer. The sizes spanned from 3 pixels to 11 in strides of 2. The number of kernels followed the powers of 2 and were varied from 2 to 64. In the case of N=1, this represented 30 different CNNs with their own architecture hyperparameters. Fig4A shows the performance (MAE averaged over three original targets and averaged over 5 folds) of each of these 30 models.

**Fig4.**
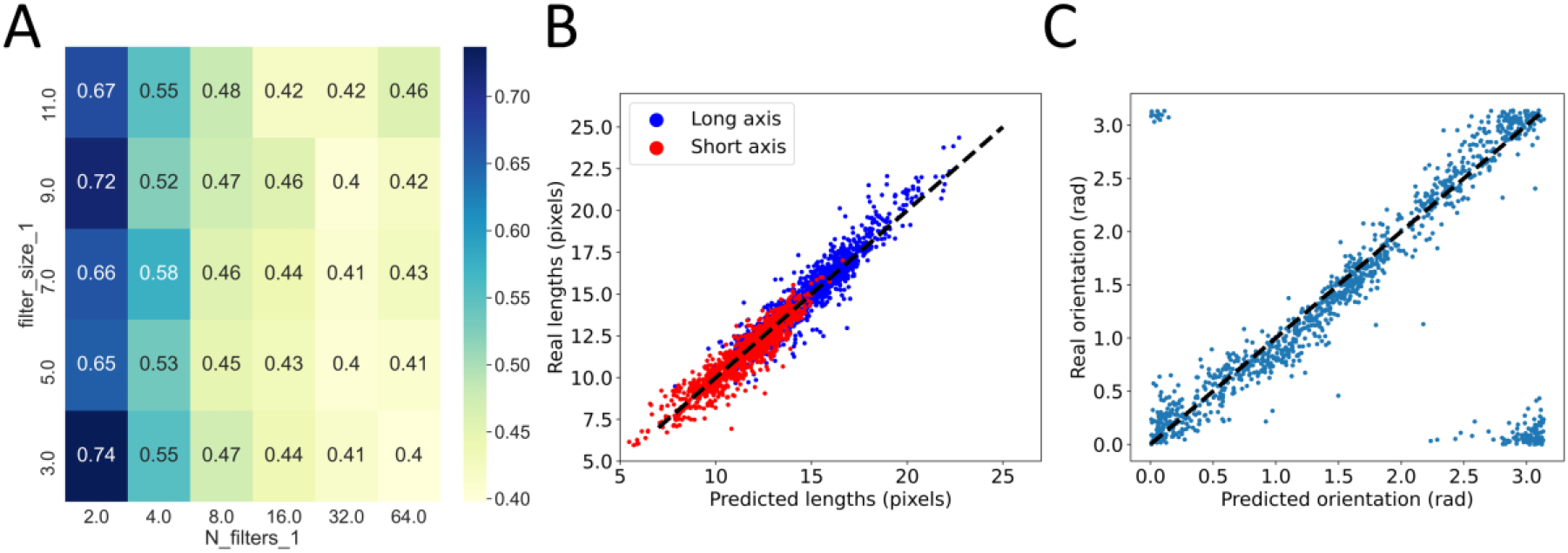
Grid search on 1-conv models. A: the MAE averaged over 5 folds is shown both in color and numbers as a function of the two varying parameters, number of filters in the convolutional layer (N_filters_1) and size of these filters (filter_size_1). The numbers correspond to the average MAEs on the three original targets. B: comparison between ground truth and model predictions for the optimal 1-conv model after retraining. Comparison is done both for the long axis, in blue, and the short axis, in red. C: similar comparison for orientation after inverse transformation from cosine and sine. Note that the two groups of point in the upper left and lower right corners are a signature of the periodicity of this measure and do not indicate bad predictions. In B and C, black dashed lines are a guide for the eye representing identity.

From this, we noticed that the number of filters seemed to have a much larger impact on performance than the actual size of these filters. However, we could also observe that above a threshold, here 16, increasing the number of filters no longer improved the performance of the model. We confirmed these results by looking separately at the effect of both parameters (Fig SI4).

In addition to these rational rules of constructions, we also extracted from this grid search the best performing model which, here, had 64 filters of size 3. This selection was based on the 5-fold cross validation and we then retrained this model once with the entire training set (except a small validation set meant to stop training in case of overfitting) and looked at its ability to generalize to the test set. Comparison between the predictions of the model and the ground truth are shown for the long and short axes (Fig4B) and for the orientation after the inverse transformation from cosine and sine (Fig4C). In both cases, we found a strong correlation between predictions and ground truth demonstrating that even this extremely simple CNN was capable of extracting valuable information. Quantitatively, this model achieved a MAE of 0.60 pixels on the long axis, 0.51 pixels on the short axis and 0.14 radians on the orientation (Table1). Still, because of its simplicity, this model was far from being optimized and we then moved on to N=2, adding a second convolutional layer, and asked whether this change could improve performance.

**Table1.** Comparison of all four final models in terms of performance on test set (MAEs), complexity, number of trainable parameters and training and analysis time. a.u. stands for arbitrary units since all times are dependent on hardware configurations.

| **Models** | **1-conv** | **2-conv** | **3-conv** | **transfer** |
| --- | --- | --- | --- | --- |
| Long axis MAE (pixels) | 0,6 | 0,31 | 0,36 | 0,44 |
| Short axis MAE (pixels) | 0,51 | 0,2 | 0,19 | 0,32 |
| Orientation MAE (rad) | 0,14 | 0,07 | 0,06 | 0,16 |
| Trainable parameters | 1049220 | 1100132 | 246676 | 1049220 |
| Total parameters | 1049220 | 1100132 | 246676 | 21073604 |
| Time per epoch (a. u.) | 132 | 306 | 255 | 1560 |
| Time to analyse 1000 images (a.u.) | 2.57 | 5.55 | 5.62 | 86.1 |

### 4 parameter grid search on 2 convolutional layer models

We repeated the exact same approach except that by adding a second convolutional layer, we now had four parameters that were varied independently over the same ranges as before. This led to a total of 900 different models that we trained using a 5-fold cross validation scheme. Note that the vast majority of these models didn’t obey the usual construction rules. The number of filters could even decrease from the first to the second layer and the same applied to filter sizes.

First, we studied the impact of filter sizes by pooling all models sharing the same filter sizes in both layers (Fig 5A). In the range of values explored, we found a limited impact of filter sizes. Still, it appeared that larger filters slightly increased performance as long as sizes were large enough in both layers (Fig 5A). A rule of thumb that emerged to tackle our problem was thus that one should use kernels of size 7 or more in both layers but that the exact sizes had limited impact (Fig 5A).

**Fig5.**
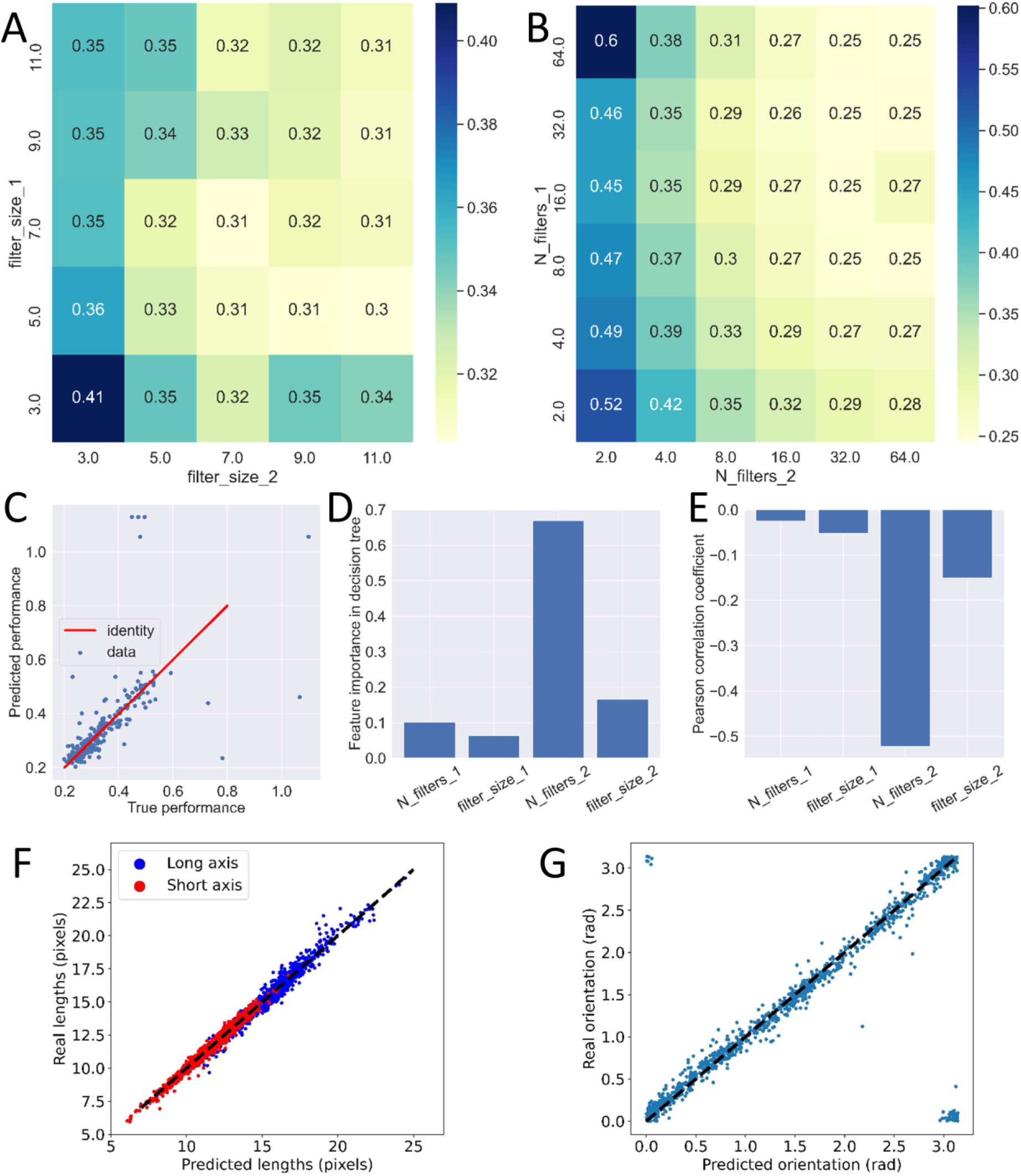
Grid search on 2-conv models. A: Effect of filter size in each layer. The averaged MAE over 5-folds of all models sharing the same filter sizes are again averaged together to yield a single MAE, shown in colors and numbers. B: same study for the effect of number of filters in both layers. Numbers in A and B represent the same quantity as in Fig 4A. C: comparison between true performance of 270 2-conv models and the performance predicted by a decision tree regressor. The red line is a guide for the eye representing identity. D: feature importance of all four parameters in the decision tree regressor showing the particular importance of N_filters_2, the number of filters in the second convolutional layer. E: Pearson correlation coefficient of all parameters with the performance of the models. Negative values indicate that increasing any of these parameters will tend to reduce the MAE and hence increase performance. F: comparison of predicted and true long and short axes for the re-trained optimal 2-conv model. G: comparison of predicted and true orientation for the same model. In F and G, the dashed black line represents identity.

Second, we used the same approach to explore the effect of the number of kernels (Fig 5B). Here, we found that a low number of kernels in the second convolutional layer had a clear negative effect on performance (Fig 5B). On the opposite, a small number of kernels in the first layer could be rescued by increasing the number in the second layer (Fig 5B). For instance, models with two and sixty-four kernels in the first and second layers, respectively, achieved performances similar to the best solutions. We also found, similar to kernel sizes, that if each layer had enough kernels (16 or more), the resulting performance was not significantly impacted by the choice of architecture.

However, these results are based on pooling together models which can have very different performances and focusing on the average might be misleading. Therefore, we also took another approach. We focused on models that follow the construction rules usually applied, e.g. keeping the same kernel sizes in both layers. This allowed us to look at the effect of the number of kernels in an unbiased way. The qualitative results found on pooled models remain true with this approach (Fig SI5). Similarly, we looked at models following the rule that one should double the number of kernels between the first and second layers and focused on the effect of kernel sizes on these models. Here too, we found that the qualitative observations from pooled models also hold true for single ones (Fig SI6).

Finally, we tried to rationalize these emerging rules by asking whether we could predict the performance of a given model from its architecture details. We therefore defined a new machine learning problem. The inputs were the four numbers representing the model architecture and the target variable was the average (over 5 folds) MAE of the resulting model. We created a dataset for this problem from our 900 pre-trained models and split it into training and test sets in order to train a regressor decision tree. We found that this simple model was quite accurate at predicting model performances (Fig 5C). Thanks to its simplicity, a decision tree is an interpretable model and the importance of each input feature can be measured. This tree confirmed what our qualitative exploration hinted at: the most important variable of all was the number of kernels in the second convolutional layer (Fig 5D). This was also confirmed by calculating the Pearson correlation coefficient of all four variables with the target (Fig 5E).

In terms of comparing these models with the single convolution layer ones, we first noticed an improvement in performance in most cases as can be seen by comparing average MAE between Fig 4A and Fig 5A and B. This was in line with the usual approach of adding layers to increase the complexity of the model and thus its ability to learn complex representations. From this extensive search, we could extract the best performing model of the 900 that were trained. Interestingly, this model didn’t follow the usual construction rules as it had 16 filters of size 9 in the first layer and 64 filters of size 7 in the second one. Once properly retrained, we could test its ability to generalize to the test set and found even better correlations between predictions and ground truth than for the best single layer model (Fig 5FG). Quantitatively, this two-layer model achieved a MAE of 0.31 pixels on the long axis, 0.20 pixels on the short axis and 0.07 radian on the orientation (Table 1), a clear improvement in performance. This led us to take our CNN’s architecture one step forward and turn our attention to the case where N=3 repetitions.

### Hyperband optimization of 3 convolutional layer models

Adding a third convolutional layer put the number of models to be trained on the order of tens of thousands. At that point, a thorough grid search with full training of all these models became out of reach or, at the very least, a waste of resources. Instead, we used a hyperband tuner to explore this large parameter space.

The hyperband is an hyperparameter optimization method (38), designed to efficiently explore a parameter space with a fixed computational budget (for instance a number of training epochs). Hyperband relies on the Successive Halving algorithms, which for a given budget *B* allocates *B/n* to *n* randomly selected configurations, trains each one of them and only keeps the best half. This step is then repeated with the *n/2* remaining configuration, allocating *2B/n* to each parameter choice.

Recursively doing this allocates exponentially more budget to the best models. The issue with this algorithm is that the user has to choose between testing a lot of configurations (large *n*, small *B/n*), or having more budget for each one (small *n*, large *B/n*). This choice depends on information specific to each problem and generally unknown to the user (such as the distribution of terminal losses in the parameter space). Hyperband tackles this problem by performing a grid search on the possible values of *n*, which enables it to have for only parameter the maximum amount of resources to be allocated to a single hyperparameter configuration. Therefore, hyperband is way more flexible as it is not impacted by whether one should favor *n* over *B/n*.

Based on our results from one and two-layer models, we also decided to change the parameter space to remove the smallest numbers of filters which clearly impacted performance and allow larger numbers for all layers. We thus varied the number of filters from 8 to 128, still following powers of 2, and kept the filter sizes similar, spanning from 3 to 11. This limited the total number of models to 15,625 while keeping the most relevant values of the parameters.

We ran HyperBand on our data and tested how our models could generalize with a 5-Fold Cross Validation. The budget allocated to HyperBand was fixed by the max number of epochs it can run on a single configuration, which we set to 40 epochs as we noticed it was enough to completely train most of our models. The best model structure was the following: 8, 64, and 128 filters with sizes of 5,9 and 3, respectively.

Because of the very nature of the Hyperband optimization, we couldn’t conduct a thorough study of the effect of all parameters as we did previously. Still, we found that the best performing model didn’t follow the usual construction rules. Instead, we found it to follow rules similar to the ones we had already inferred: the size of the filters had to be large in the first layers and the number of filters had to be large in the final layers. Once retrained on the entire training set, we could quantify the ability of this model to generalize (Fig 6A-B).

**Fig6.**
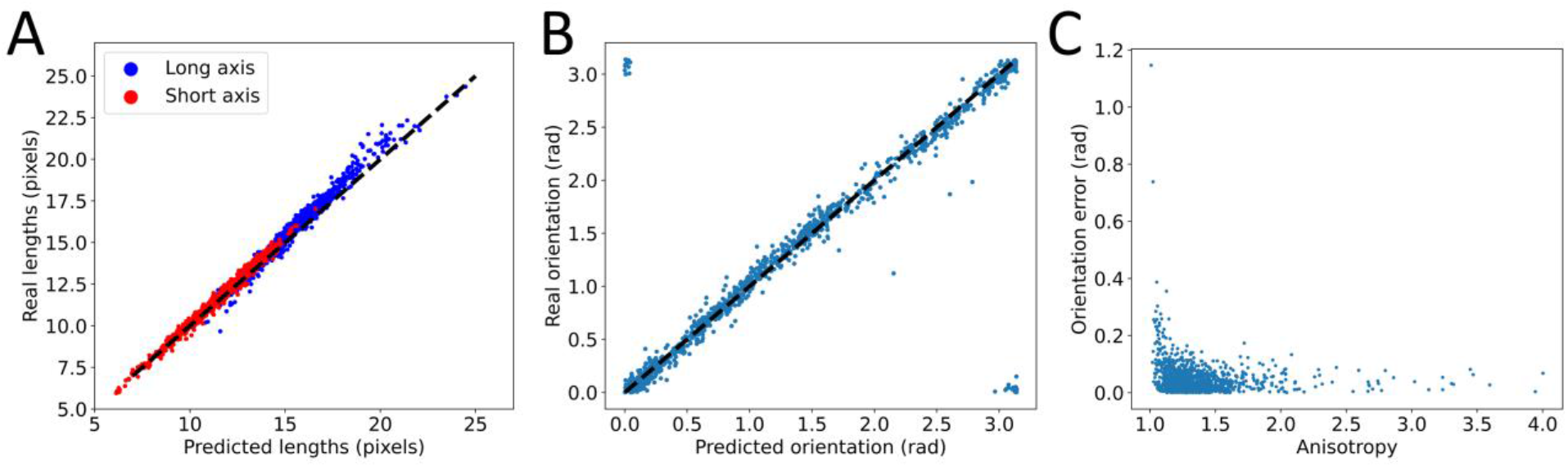
Performance of the best 3-conv model. A: comparison of predicted and true long and short axes for the re-trained optimal 3-conv model. B: comparison of predicted and true orientation for the same model. In A and B, dashed black lines represent identity. C: effect of anisotropy on the performance of the model in predicting orientation showing that the lower the anisotropy the worse the prediction.

We found the performance of this model to be very close to that of the best two-layer model with MAEs of 0.36 pixels on the long axis, 0.19 pixels on the short axis and 0.06 radian on the orientation (Table 1). Regarding orientation, we also explored another interesting property. As this orientation is that of an averaged cell over the field of view, it is strongly linked to the anisotropy of this cell, defined as the ratio of the long axis to the short axis. When anisotropy is high, there is a clear orientation of the cell and this measurement is well-defined. As anisotropy approaches one, the average cell becomes more symmetrical and orientation becomes an ill-defined quantity. We therefore explored if the performance of the model on orientation was linked to anisotropy and indeed found that this performance degraded as cells became more round (Fig 6C). This is not to be interpreted as a failure of the model but as a default in the very definition of the problem, illustrating once again the importance of problem definition and data preparation in machine learning.

Overall, this 3-layer convolutional network showed only marginal, if any, improvement when compared to the best 2-layer one. This was unexpected but it was already performing well enough to be a relevant tool to measure cell shapes on a coarse grained level. In terms of long and short axis, the MAEs we obtained on the test set corresponded to an average relative error of 2.4% and 1.7%, respectively, which is an improvement from previous methods (20) and clearly acceptable as a measuring tool.

For all these reasons, we decided to stop at this step and consider we had successfully solved the image analysis problem we defined by designing simple, optimized CNNs. Our approach revealed the important parameters to optimize and showed that adding convolutional layers improved performance up to a point. Researchers faced with very similar images to the ones we used here could directly re-use these models to extract cell shape information at a coarse-grained level from confluent tissues. However, the results obtained here and the ability of our models to generalize to other situations highly depends on the dataset used for training. It is a common feature in deep learning that models will fail when facing different distributions of the inputs, targets or both. We argue here that researchers in such situation should build and train their own simple CNNs rather than transferring from other models, even the ones presented here. To do so, a large annotated dataset is necessary and is often the limiting factor to conduct a successful machine learning approach. This is the reason why transfer learning is widely used as it promises good performance with a smaller dataset and less technical knowledge since the models themselves don’t have to be rebuilt from scratch. We thus tested this transfer approach and compared it to our own.

### Transfer learning from pre-trained models

Many pre-trained models are easily available in dedicated libraries. Most of the time, people use complex models that have been designed and trained for the ImageNet dataset and competition: the ImageNet Large Scale Visual Recognition Challenge. This is probably the gold standard in computer vision and has naturally attracted most of the novelties in the field. For example, this is where both the Visual Geometry Group (VGG) (39) and Residual Networks (ResNet) (40) models were first introduced and these remain the models of choice for transfer learning. Here, we didn’t perform a thorough investigation of all available models and focused on one of them: VGG-19. This model has 16 convolutional layers, 5 maximum pooling, 3 dense layers and over 140 million trainable weights. As a basis for comparison, the most complex of our optimized model has a little over one million trainable weights (Table 1). VGG19 has frequently been used in transfer learning approaches including in biomedical research (41–43).

To adapt this model to our problem, several steps had to be carefully followed. First, we loaded and froze all convolutional layers of this model and removed its dense layers at the top. We replaced them with our own dense layers, one with 128 neurons and Relu activation and our final regressor with 4 neurons with linear activation. This architecture was chosen to give this model the same order of trainable parameters, around one million, to have a direct comparison with our own CNNs and reduced the total number of parameters to a little over 20 million (Table 1). Second, since the first layers of the model were frozen, we needed to adapt our inputs to resemble those on which VGG19 was trained. This meant changing the size of the input layer to fit our 128*128 images and artificially turning them into color images with three different channels whereas our raw images were black and white. Finally, we used the same learning strategy as before, again to have a direct comparison. We argue that all these steps, in addition to problem definition and creation of a proper data set, makes transfer learning as, if not more, technically difficult as designing one’s own models.

With these changes implemented, we then trained this new model (more precisely its top dense layers) on the same training set as before and tested its performance on the test set. We found that this complex model didn’t improve performance (Fig 7A) and achieved MAEs of 0.44 pixels on the long axis, 0.32 pixels on the short axis and 0.16 radians on the orientation. Comparing with our three other CNNs (Fig7 B-D), we find VGG19 to perform better than our best 1-conv model but slightly worse than the 2-conv and 3-conv optimized models. Overall, it seemed clear that this approach did not yield better performances.

**Fig7.**
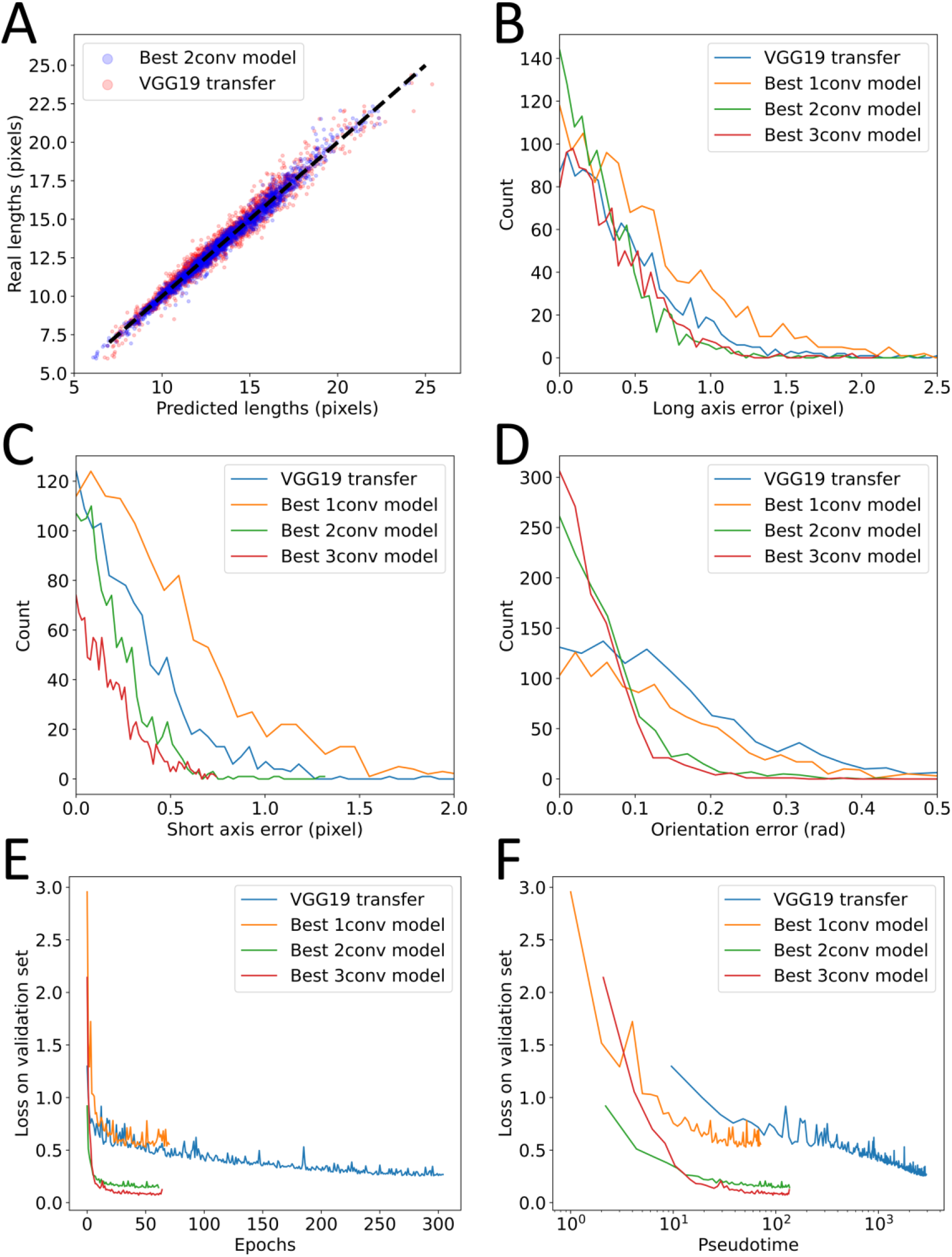
Transfer learning. A: Comparison of true and predicted long and short axes for the best 2-conv model (blue) and transfer from VGG19 (red). The wider scatter of red dots indicates lower performances. The black dashed line represents identity. B-C-D: distribution of absolute errors on the test set for all four models and for long axes (B), short axes (C) and orientation (D). E: Loss on validation set during the final training of each model as a function of the number of epochs. Note that training is automatically stopped when no improvement is achieved for 20 epochs, explaining the different lengths of the plots. F: Loss on validation set as a function of time spent training. Note that time is normalized since real training times would be dependant on the computer used for training.

At some point, marginal improvements on performance are no longer a priority and the question of computing time and resources becomes more relevant. We thus decided to compare the efficiency of our different strategies in terms of training time. In that respect, small homemade CNNs also outperformed VGG19. Not only did VGG19 require more epochs to converge (Fig 7E) but each epoch also took longer to compute. Although they all have comparable numbers of trainable parameters, thus requiring similar number of gradient descent and weight updating calculations, our modified VGG19 had over 20 million non-trainable parameters. This implies that the treatment of each individual image required more calculation which made each epoch take longer. Similarly, after training, the time required to analyze one image was significantly longer for VGG19 than for any of our CNNs. All combined, we found that VGG19 took ten times longer to fully train than any of our simple models (Fig 7F, Table 1) while achieving lower performances.

There is one more argument usually put forward in favor of transfer learning: it requires fewer annotated data to train properly. This is a very important argument since large, high quality datasets are difficult to build and are the main added value in most machine learning applications. Again, we were able to conduct a thorough investigation of CNN architecture rules thanks to our large, well annotated dataset but we acknowledge that this is more the exception than the rule.

To test this argument in our context, we decided to artificially limit the training set and monitor the performance of our optimized 3-conv model and VGG19 as a function of the number of images accessible in this training set. We found that even with very low number of images, down to a few tens, the performance of VGG19 was not significantly better than our homemade 3-conv CNN (Fig 8) for all of our different targets. Finally, we found, as expected, that after training, the time required to analyze new images was longer for this more complex model (Table1).

**Fig8.**
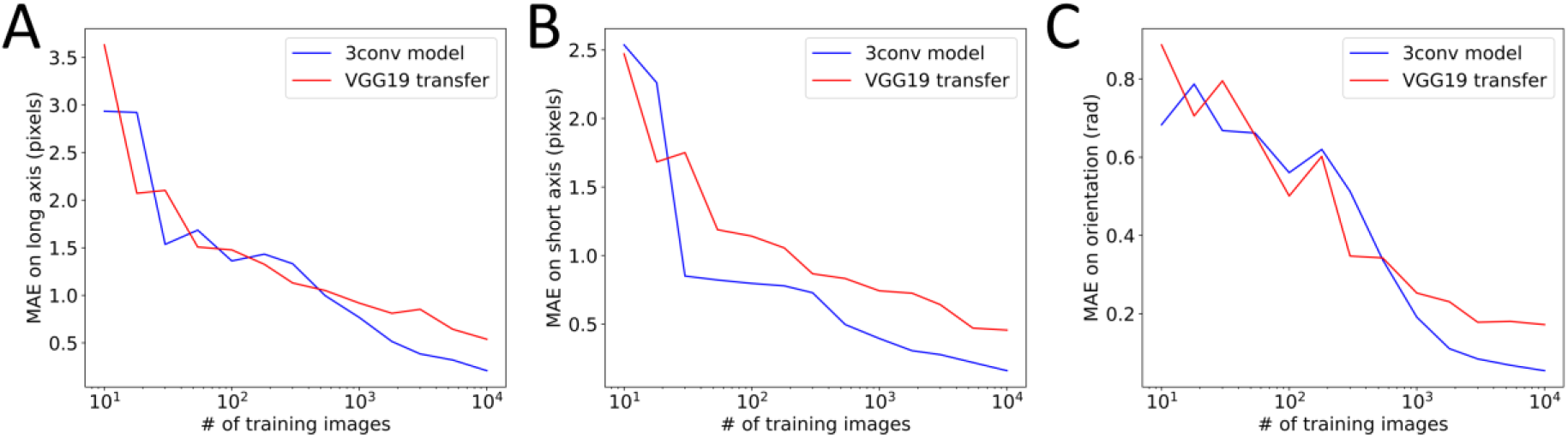
Effect of training set size. Performance of either the best 3-conv model (blue) or transfer from VGG19 (red) as a function of the number of images used in their training. MAE is shown for long axis (A), short axis (B) and orientation (C). No clear trend appears to claim that one model is more performant than the other.

This shows that a transfer learning approach from a complex model offers little benefit for the problem we have defined here.

## Discussion

We defined an image analysis problem relevant for the community studying the mechanobiology of living tissues at a coarse grained scale. Thanks to a pre-existing large dataset of segmentation, we defined a learning strategy, for example showing how the to deal with a difficult target such as orientation by replacing it by a couple of better defined targets, the sine and cosine of double this orientation. We then explored the performance of simple CNNs comprising one, two or three convolutional layers. We realized a thorough investigation of the impact of the network’s architecture, in the form of the sizes and numbers of convolutional filters and found that the optimal construction rules are as follows. The number of filters in the last layers is the most impactful parameter but performance stopped increasing after this number passed a threshold. The sizes of the filters seemed to have a limited influence on performance as long as they were large enough, they could even be varied from one layer to the next. In the end, we settled on an optimized 3 convolutional layer model which demonstrated a high capacity to generalize to new data with an average relative error of a few percent. This constitutes our first important result in the form of an efficient way to measure cellular shapes in a confluent tissue without the use of segmentation and all the drawbacks it entails.

Still, our approach was made possible by the large amount of high quality data and we acknowledge that this is not a common situation in practice. We thus explored the possibility to use transfer learning from VGG19 to address the same problem. Interestingly, we found that this approach required as much, if not more, skills in coding and machine learning, that it yielded performances barely on par with our simpler CNNs, that it required ten times longer to train and analyze new data and that even with a small number of training data, it wasn’t more efficient than our simple approach.

In addition to the resulting optimized 3-layer model, our work also suggests a roadmap for researchers facing an image analysis problem well suited for CNNs but which remains simpler than the image classification of ImageNet on which most pre-trained models were optimized. Instead of using transfer learning from these models, we argue that it is simpler, faster and more efficient to build from scratch simple CNNs with only a few convolutional layers, respecting simple rules of construction. If the dataset allows, an optimization of hyperparameters such as the architecture of the models can yield great performance and even provide information on the complexity of the problem itself.

As a conclusion, we illustrate this strategy on a different, albeit very similar problem. We used images of cell flow during primitive streak formation in a chicken embryo (Fig 9A) and wanted to create maps of cellular anisotropy over these very large tissues. Thanks to semi-manual segmentation over the all image (20,44), we created a dataset of over 3,000 images each containing around ten cells (Fig 9B) and for which we had a measurement of the average cellular anisotropy.

**Fig9.**
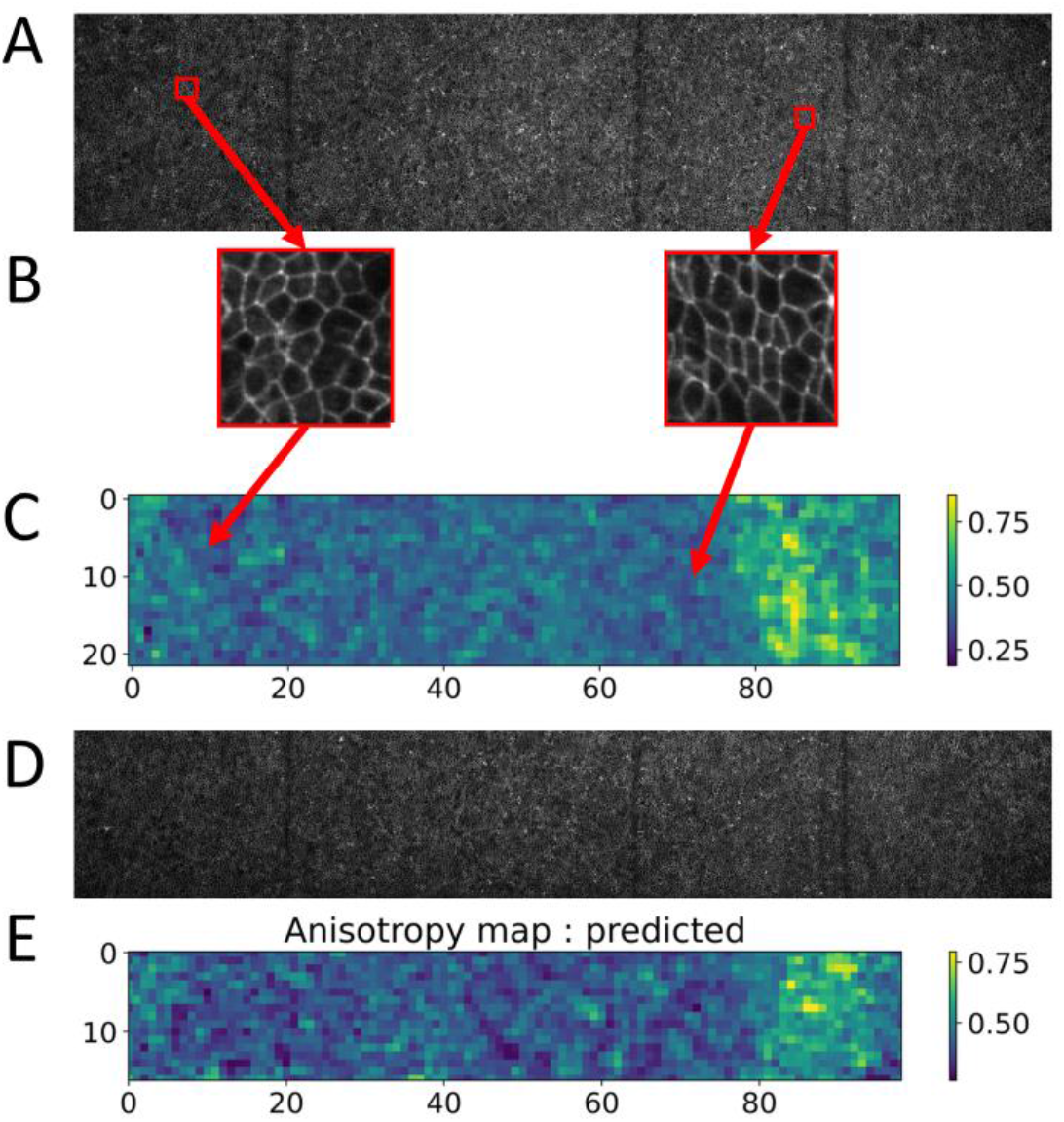
Illustration of strategy. A: Original image used to create a training set. It is the top part of a larger image which was separated in two. B: two examples of 128*128 images extracted from A. C: anisotropy map of the full image, each image in B represents one pixel of this map. D: Original image used to create a test set which was separated in similar 128*128 windows and these images were passed to the trained CNN. E: Predicted anisotropy map of the image in D. Comparing the ground truth in C and predicted anisotropy maps in E, we note that they both have a region of higher anisotropy visible on the right.

Each of these images will correspond to a single measurement of cellular anisotropy and thus represent one pixel of our final anisotropy map (Fig 9C). As an illustration, we used the top part of the original image to create a training set and treated the bottom of the image as new, unseen data.

Following our conclusions, we neither performed a complete grid search on hyperparameters nor did we try to use transfer learning from complex models or our own small CNNs optimized on a different dataset. Instead, we manually built a simple, 3 layer CNN following the construction rules we obtained: all layers have 64 filters of size 5. This network was then fully trained on the top part of the image and used to predict the anisotropy map on the bottom of that image (Fig 9D-E).

Applying this strategy, we managed to prepare the dataset, train the model and compute the predicted anisotropy map in less than a day on a laptop computer. The resulting anisotropy map (Fig 9E) shows the area of large anisotropy on the right part of the original image which is also observed on the top part of the image. This is the signature that the model correctly learned how to measure anisotropy in an efficient way.

## Material and methods

### Dataset and training strategy

The dataset is the same as the one presented in (20) and obtained in (37) and we used the same definition of the average long axis, short axis and orientation in each picture. We tranformed the orientation defined between 0 and pi as the sine and cosine of double this orientation, assuring they were both bound between -1 and 1. Major and minor axis were both normalized using a standard scaler fitted on the training set. For clarity’s sake, the output of all models were rescaled back to the original scale in pixels.

The images were normalized by 255, the maximum value in 8-bit images, both in the training and test sets. All models were coded and trained in Python using the Keras library and Tensorflow.

For the 1-conv and 2-conv grid search on hyperparameters, each individual model was trained using a 5-fold cross validation scheme. Each training was performed using an Adam optimizer with 0.001 learning rate and with the mean squared error as loss function. This loss was monitored at each epoch on the validation set and training was automatically stopped after 20 epochs without improvement on the validation loss. We used the mean absolute error, averaged over the 5 folds and over the 4 targets, as a way to compare the performances of each model.

Once a model was selected thanks to this scheme, it was re-trained once from scratch and on the entire training set. A small part (10%) was kept as a validation set to be used for early stopping with the same criteria as above. The model then predicted the test set, which had remained unseen up until now, and the measure of performance on this test set were used throughout this paper as a measure of their ability to generalize to new data. For Fig8, a subset of the training set was randomly selected to perform the training of both the optimized 3-conv model and transfer from VGG19 and the resulting performances were measured on the entire test set.

For clarity and visualization purposes, we also did the inverse transformation from cosine and sine back to the actual orientation by taking the 2-argument arctangent and dividing it by 2.

### Comparison of training strategies

To define and test our training strategy, we trained 2 convolutional layers CNN respecting the architecture shown in Fig 3 with 32 kernels of size 5 in each layer. The output layer was changed to match different strategies. For Fig SI1, the “mixed” model we had three output neurons to predict three targets as we left the orientation as originally defined. We then had to modify the loss function to reflect the periodic nature of that target. The final loss function for training was thus the sum of standard MSEs on long and short axes and the square of the orientation error modulo pi. The sincos model used four output neurons to predict axes and both sine and cosine of the orientation. In Fig SI2, we trained two “dedicated models”, each with two output neurons to predict either both axes or sine and cosine of orientation.

### Decision tree

We used a decision tree regressor from the scikit-learn library to predict the performance of 2-conv models from the 4 architecture parameters we varied. We kept all hyperparameters of the tree as default except for the maximum depth which was set to 16. The target was defined as the MAE on validation set averaged over the 5 folds during training of the model. This tree was trained on 630 points out of 900 and tested on the remaining 270.

We used the feature importance in this tree and a Pearson correlation coefficient as two different measures of the impact of each of the 4 parameters.

### Hyperband

We used a modified version of the Hyperband algorithm from KerasTuner module. It was modified to include cross-validation in the hyperparameters tuning, testing our models on a 5-fold split of the training set, with the mean of the Mean Squared Error across all splits as loss function. This was achieved by replacing the training line in the tuner’s code by a loop on the 5 splits, training 5 different models instead of one at each step of Hyperband. Training parameters are identical to the one used in grid search.

We then ran our modified Hyperband on 3-conv models, allowing the number of filters on each convolution layer to be any power of 2 between 8 and 128, with kernel size ranging from 3 to 11 by steps of 2 and a maximum pooling of size 2. We set the maximum number of epochs to 40, as we noticed it was enough for any model we encountered to be fully trained.

The last parameter to be fixed is the number of Hyperband iteration the user wants to run. We recommend setting this as high as one can afford computation-wise. KerasTuner provides ways to save the best model, therefore HyperBand can be stopped before completion if the performance are satisfactory.

### Transfer

We loaded the VGG19 pre-trained model from Keras and used the corresponding ImageNet weights. We modified the input layer to fit the size of our images of 128*128 pixels. We didn’t load the top part of the model and froze all remaining layers. We then manually added two consecutive dense layers with, respectively 128 neurons with Relu activation and 4 neurons with linear activation. The targets were treated the same way as for our own CNNs.

In terms of inputs, we pre-treated the images in both the training and test sets using the preprocessing function provided especially for VGG19. The model was then trained once with the same optimizer, learning rate, early stopping and batch size as our own CNNs. For example, here too, 10% of the training set was kept as validation to monitor early stopping.

### Chicken

The data set was obtained in (44) and later used in (20). It consisted in a single large image of a chicken embryo during primitive streak formation. The entire image was accompanied by a semi-manual segmentation used to define the ground truth. This original image was split in the middle, the upper part being kept as a training set and the bottom one as a test set. Each set was cut in 128×128 pixels images with 25% overlap and the average long and short axes were computed from the segmentation for each resulting image. This yielded over 3,000 images in the data set.

We designed a three layer CNN with the architecture shown in Fig 3 with the addition of a dropout layer after each convolutional one with a dropout of 0.2, and with only two output neurons to predict long and short axes. The model was trained once on the training set, keeping 20% of it as a validation set to apply early stopping and avoid overfitting. Once trained, the model was then used to predict these two quantities on the test set to predict local anisotropy.

Anisotropy was defined as 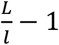 with *L* and *l* the long and short axes, respectively.

### Sine-cosine learning

The relationship between cosine and sine learned by the model and presented in Fig SI3 was obtained by the optimized 2-conv model after full retraining. It shows the relationship between predicted cosines and sines on the test set.

## Acknowledgments

We thank Yohannes Bellaiche and Cornelis Weijer for the fly thorax and chicken embryo images and data sets. We also acknowledge help from Osvanny Ramos and Victor Levy dit Vehel for fruitful discussions in designing the project. Finally, we thank Charlotte Rivière for feedback on the manuscript.

