## Supplementary figures for "Small hand-designed convolutional neural networks outperform transfer learning in automated cell shape detection in confluent tissues"

### Supplementary figure

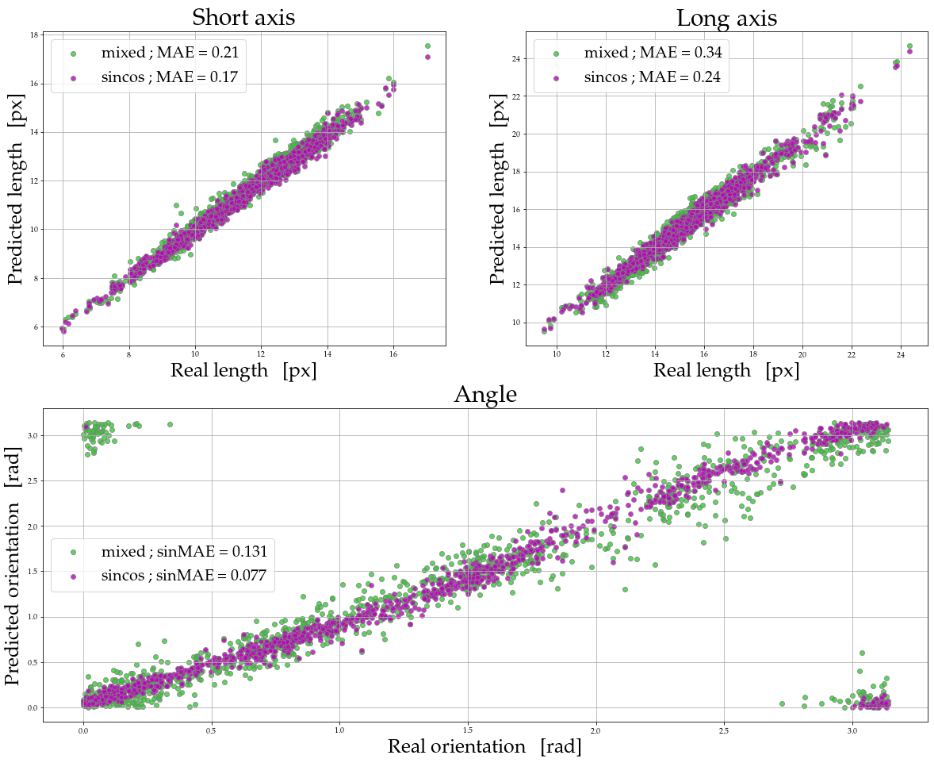


*SI1. Definition of strategy. Comparison of performance of a simple 2 convolutional layer CNN trained either with the raw orientation and a circular MSE (mixed) or by encoding orientation as sine and cosine (sincos). In each case, the corresponding MAEs are shown in inserts.*

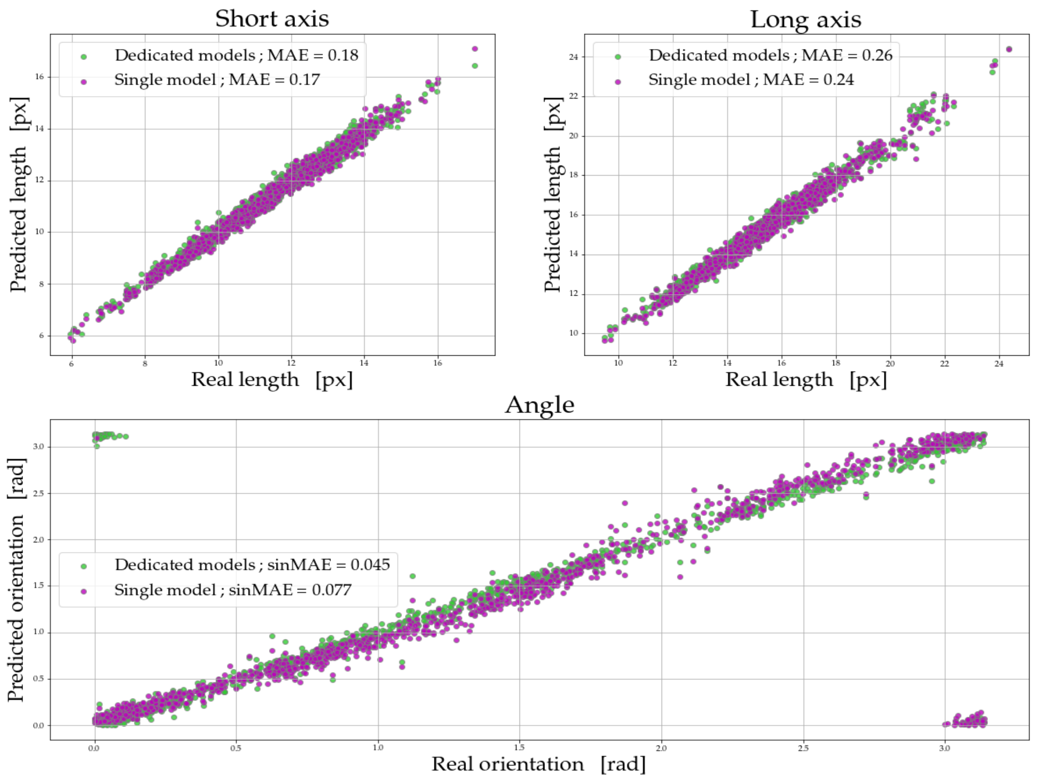


SI2. *Definition of strategy. Comparison of performance of the performance of simple 2-conv CNNs. The dedicated models are two different CNNs, one trained to predict lengths only and the other orientation only. The single model predicts all three quantities at once. In each case, the corresponding MAEs are shown in inserts.*

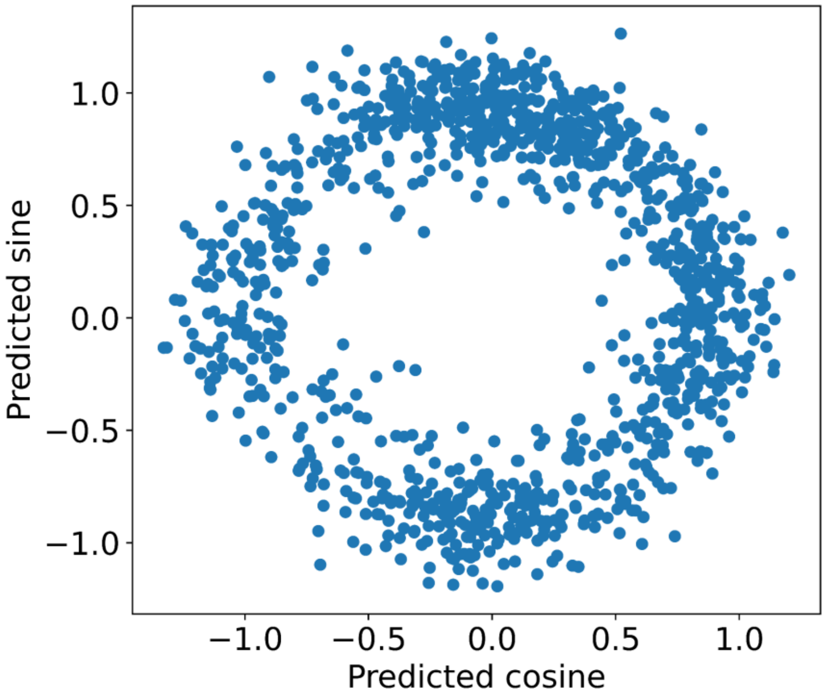


*SI3. Learning of trigonometric relations. Correlation between predicted sines and cosines from a 2-conv layer CNN on the test set.*

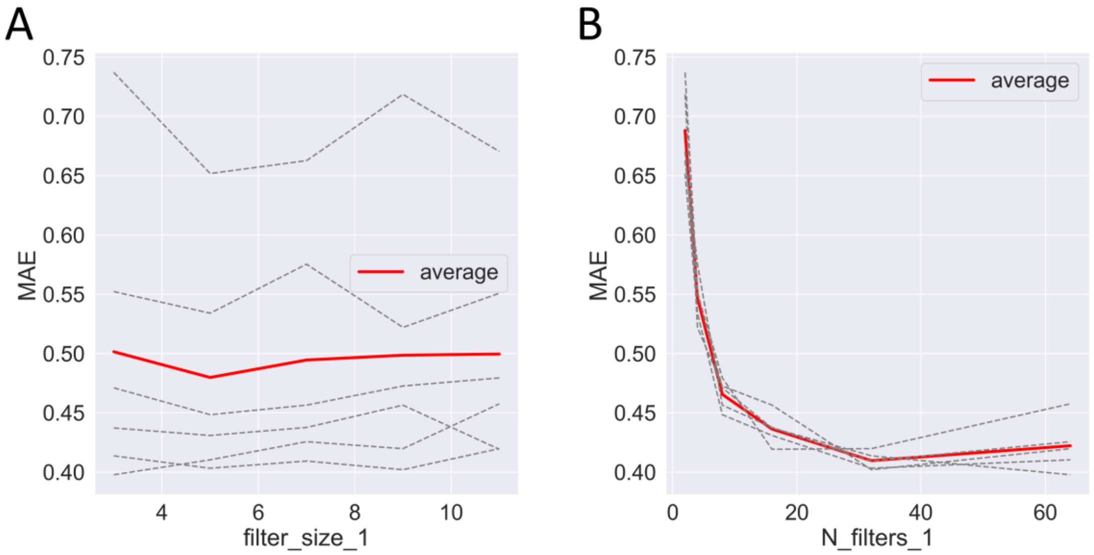


*SI4. Effect of filter size (A) and number of filters (B) on MAE over 5 folds for 1-conv models. In A, each dashed black line corresponds to one value of the number of filters and the red line is the average of all black lines. In B, each dashed black line correspond to one value of filter size and the red line is the average of all black lines.*

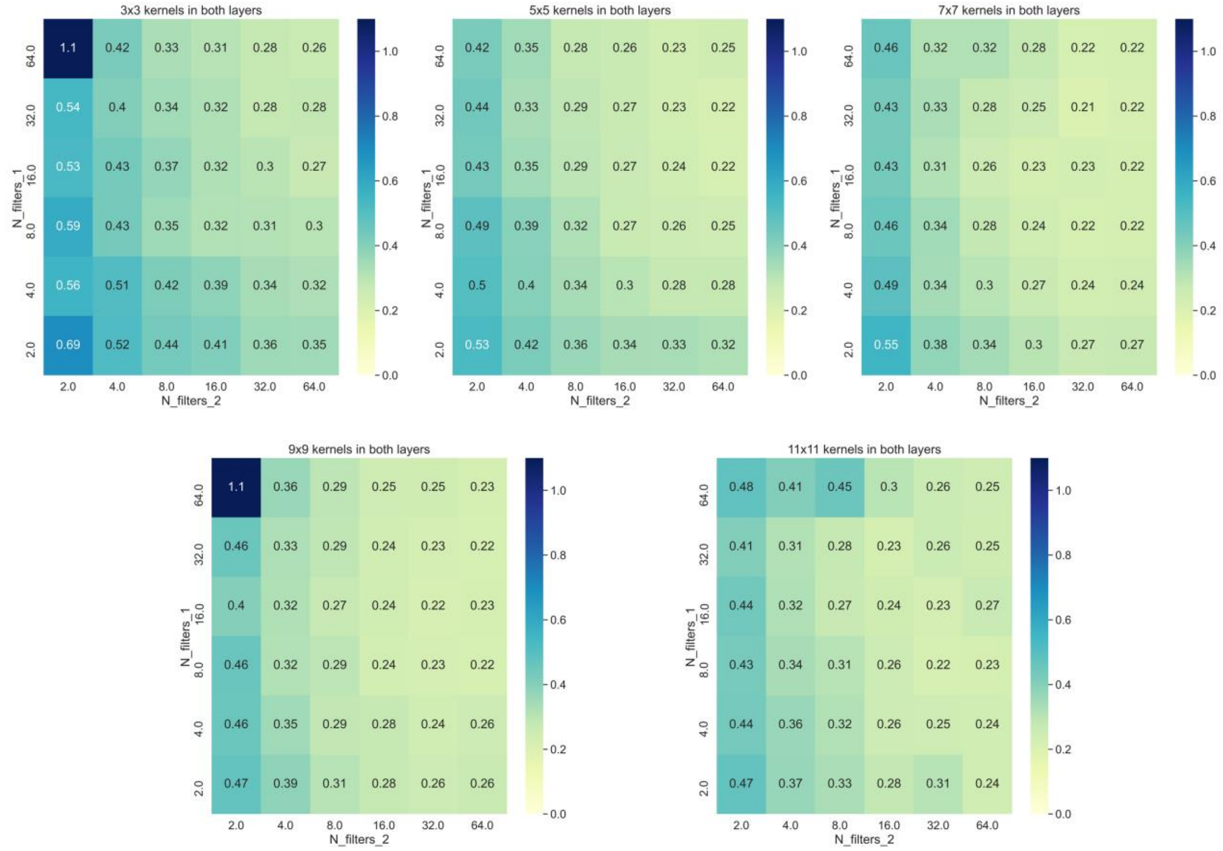


*SI5. Performance analysis of 2-conv models individually. Each graph shows models where both layers have the same filter sizes (from 3 to 11). Each value of the MAE averaged over 5-fold is shown both in colors and numbers and corresponds here to a single network architecture, or a single model.*

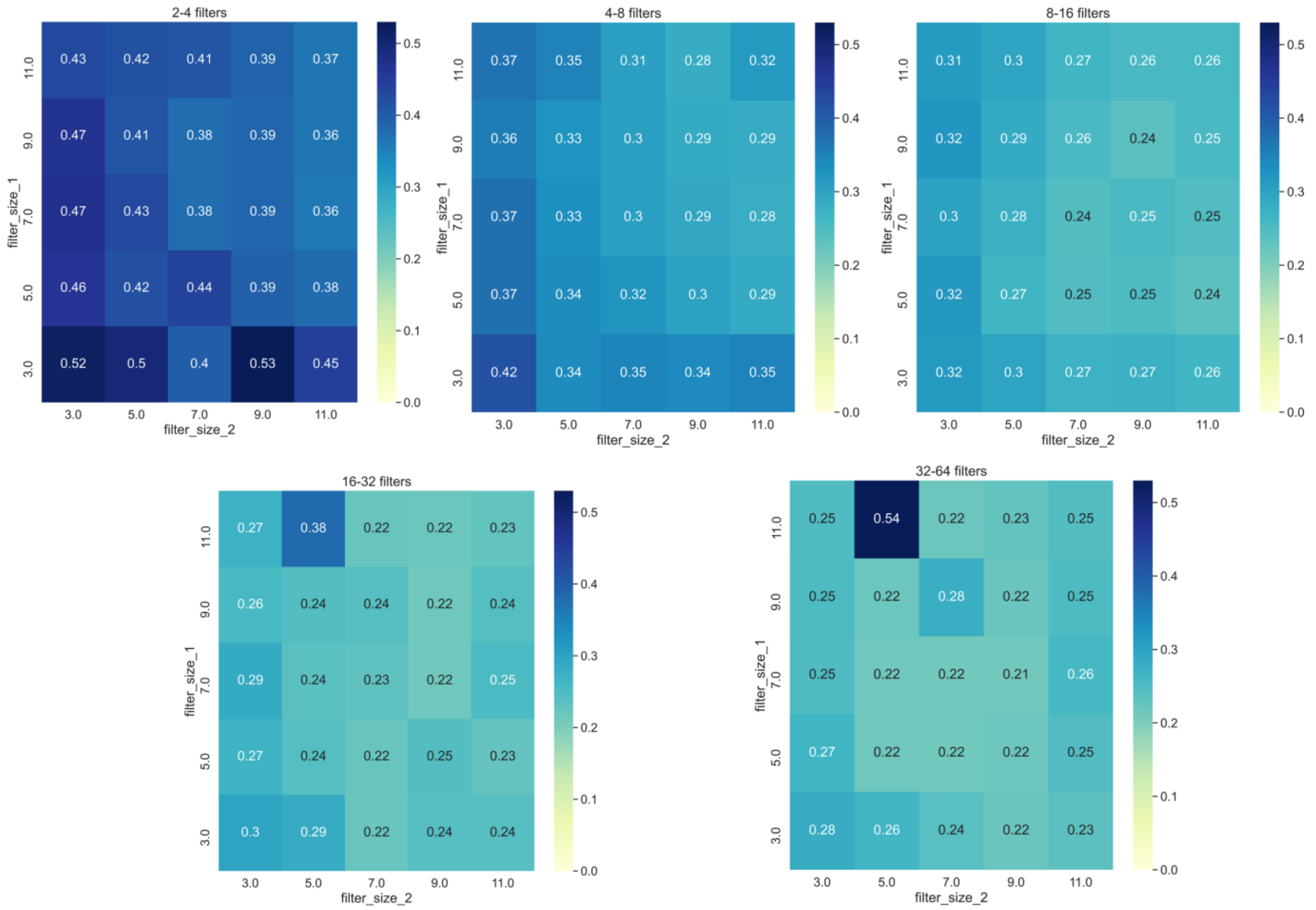


*SI6. Same analysis as Fig SI5 but for the sizes of the filters. Each model in a graph respects the rule that the number of filters is multiplied by 2 between the two layers and these values span from 2-4 to 32-64.*
